# Single-cell tracking reveals super-spreading cells with high persistence in invasive brain cancer

**DOI:** 10.1101/2020.10.06.327676

**Authors:** Aimilia Nousi, Maria Tangen Søgaard, Liselotte Jauffred

**Affiliations:** The Niels Bohr Institute, University of Copenhagen, Blegdamsvej 17, DK-2100 Copenhagen O, Denmark

## Abstract

Cell migration is a fundamental characteristic of vital processes such as tissue morphogenesis, wound healing and immune cell homing to lymph nodes and inflamed or infected sites. Therefore, various brain defect diseases, chronic inflammatory diseases as well as tumor formation and metastasis are associated with aberrant or absent cell migration. With embedment of multicellular brain cancer spheroids in Matrigel™ and single-particle tracking, we extracted the paths of cells migrating away from the spheroids. We found that - in contrast to local invasion - single cell migration is independent of the mechanical load exerted by the environment and is characterized by high directionality and persistence. Furthermore, we identified a subpopulation of super-spreading cells with >200-fold longer persistence times than the majority of cells. These results highlight yet another aspect of between-cell heterogeneity in tumors.

## INTRODUCTION

Tumor cell migration is a hallmark of cancer and is typically divided into three major types: amoeboid-, mesenchymal-, or collective migration Hanahan and Weinberg 2011. Amoeboid and mesenchymal migration is the movement of single cells. Although cancers tend to exhibit one specific kind of *migration* - such as amoeboid migration by lymphoma cells - migrating cancer cells are known for their high plasticity and ability to interconvert between different modes of migration in response to environmental cues Friedl and Wolf 2003, Friedl et al. 2012. In contrast, collective migration i.e. *invasion*, is the cooperative transport of whole groups of cells, which describes the local expansion of tumor cells into the extracellular matrix (ECM).

A common system used to model tumors in 3D is that of tumor spheroids. Spheroids are dense multicellular structures that form due to most cells’ natural tendency to aggregate under certain environmental conditions Anton et al. 2015. The advantages of spheroids as tumor models, include minimal interface between cells in culture and the local environment, increased cell-cell adhesion and tight junction formation; as well as the existence of nutrient, oxygen and cell proliferation gradients from the surface to the core Phung et al. 2011, Yamada and Cukierman 2007. Cells in spheroids therefore exhibit different phenotypes depending on their proximity to the surface much like cancer cells in solid tumors, which are e.g. dormant or necrotic at the core Yamada and Cukierman 2007, Nieto et al. 2016. For studies of metastasis, spheroids are embedded in a collagen, hydrogel or Matrigel™ matrix to mimic the natural cancer environment Correa de Sampaio et al. 2012, Smart et al. 2013, Goodman et al. 2007, Anguiano et al. 2017. Thus, this model system allows true three-dimensional migration/invasion of the cancer cells into the surrounding matrix.

Migration of tumor cells is a multistep process initiated by the epithelial to mesenchymal transition (EMT) where cells lose their apical-basal polarity and change their expression of e.g. surface adhesion molecules to allow a more migratory phenotype Huber et al. 2005. This transition is in itself a multistep process with multiple semi-stable intermediate states Nieto et al. 2016, Bidarra et al. 2016, Jolly et al. 2016. In order to migrate, mesenchymal cells have to develop pseudopods which protude from their body and form focal contacts with the ECM. Proteases expressed on the cell surface then up-concentrate at these contacts to degrade the ECM locally and carve tunnels for the cell as it moves. The actual motion is driven by actomyosin contraction within the cell followed by detachment of the trailing edge by disassembly of focal contacts Friedl and Wolf 2003. Due in part to the plasticity of cancer cells, the exact integrins, proteases and signaling molecules involved in this cascade of events depend not only on the cancer type but also on homogeneity among individual cells Friedl and Wolf 2003, Nieto et al. 2016

A number of different random walk models have been used to describe cell migration in various species Codling et al. 2008, Saxton 2007. Specifically, persistent random walk (PRW) models derived from the Ornstein-Uhlenbeck process have been very useful for describing the motility of various species from *Dictyostelium* over mouse fibroblasts to human endothelial cells Selmeczi et al. 2008, Dunn and Brown 1987, Gail and Boone 1970, Guisoni et al. 2018, Stokes et al. 1991. However, there are several examples of motile cell types that are not exhaustively modeled as simple persistent random walkers Selmeczi et al. 2005, Dieterich et al. 2008, Campos et al. 2010, Upadhyaya et al. 2001, Wu et al. 2014. In the specific case of glioblastoma invasion and migration, a sigmoidal Gompertzian model has been used to describe the growth of glioblastoma spheroids *in vitro* Chignola et al. 2000, while a cellular automaton model was used to highlight the importance of cell-cell adhesion/attraction during glioma cell migration Aubert et al. 2006. Further, the PRW model has been used to describe both the growth and diffusion of cells away from glioblastoma spheroids Stein et al. 2007. While this work separated the proliferation and migration into two distinct populations, it has been shown that the combined growth and migration can be described by a density-dependency, i.e., cell migration is inversely proportional to cell density, which is highest in the proliferating zone Khain et al. 2005, Stepien et al. 2015, Rutter 2016. A number of studies have also modeled glioblastoma behavior in regard to morphology, metabolism, vasculature, and medical treatment Martirosyan et al. 2015. However, studies on the dynamics of glioblastoma cell migration are still limited and there seems to be no consensus on the subject.

We therefore set out to further characterize the invasion and migration of brain glioblastoma cells (U87-MG) from multicellular tumor spheroids, a cell type forming particularly tight spheroids Vinci et al. 2012. We developed a brightfield live-cell imaging setup to follow the invasion of Matrigel™-embedded spheroids as well as subsequent migration of individual cells away from it. We found that the ability to invade the extracellular matrix is compromised by matrix stiffening. However, our analysis - based on machine learning algorithms Sommer et al. 2011 - also yielded insights into the migration behavior of individual cells and we found this to be highly ECM density independent. Surprisingly, we identified a subpopulation of fast migrating cancer cells. We found that these *super-spreaders* are not moving significantly faster along their trajectory, but with much more persistence.

## RESULTS

We cultured gravitation-assisted brain cancer spheroid by seeding of ∼ 650 U87-MG cells in 200 µL low-attachment U-formed wells. After 4 days of incubation, spheroids with an average diameter of 276 ± 6 µm (means ± SD, *N* = 34), an average aspect ratio of 1.34 ± 0.01 (means ± SD, *N* = 34), and a density of ∼ 3 · 10^−4^ cells/µm^3^ Niora et al. 2020 were harvested. To kick-start invasion, we interchanged growth medium with the ECM formulation Matrigel™ which consists of laminin, collagen IV, fibronectin, and various growth factors and, thus, mimics basement tissue membranes Engbring and Kleinman 2003. We used a range of concentrations from 25% to 75%, thus subjecting spheroids to various mechanical load Zaman et al. 2006 and imaged the spheroids over more than 24 hours (maximum 31 hours) with a frame rate of 1 every 5 minutes. All our data was obtained with simple wide-field microscopy, which limits sources of potential phototoxicity but complicates image segmentation. Examples of frames from the resulting time-lapse movies is shown in figure 1A for a spheroid in 50% ECM at time *t* = 0, 8, 16, 24 hours. Here it is obvious that there are two types of motility in play: i) branch-like pattern of cells moving collectively through the ECM as well as ii) individual cells leaving the spheroid body. These different motility modi are well-known Vinci et al. 2012, Berens et al. 2015, Motaln et al. 2015 and corresponds to invasion and migration, respectively.

**Figure 1:**
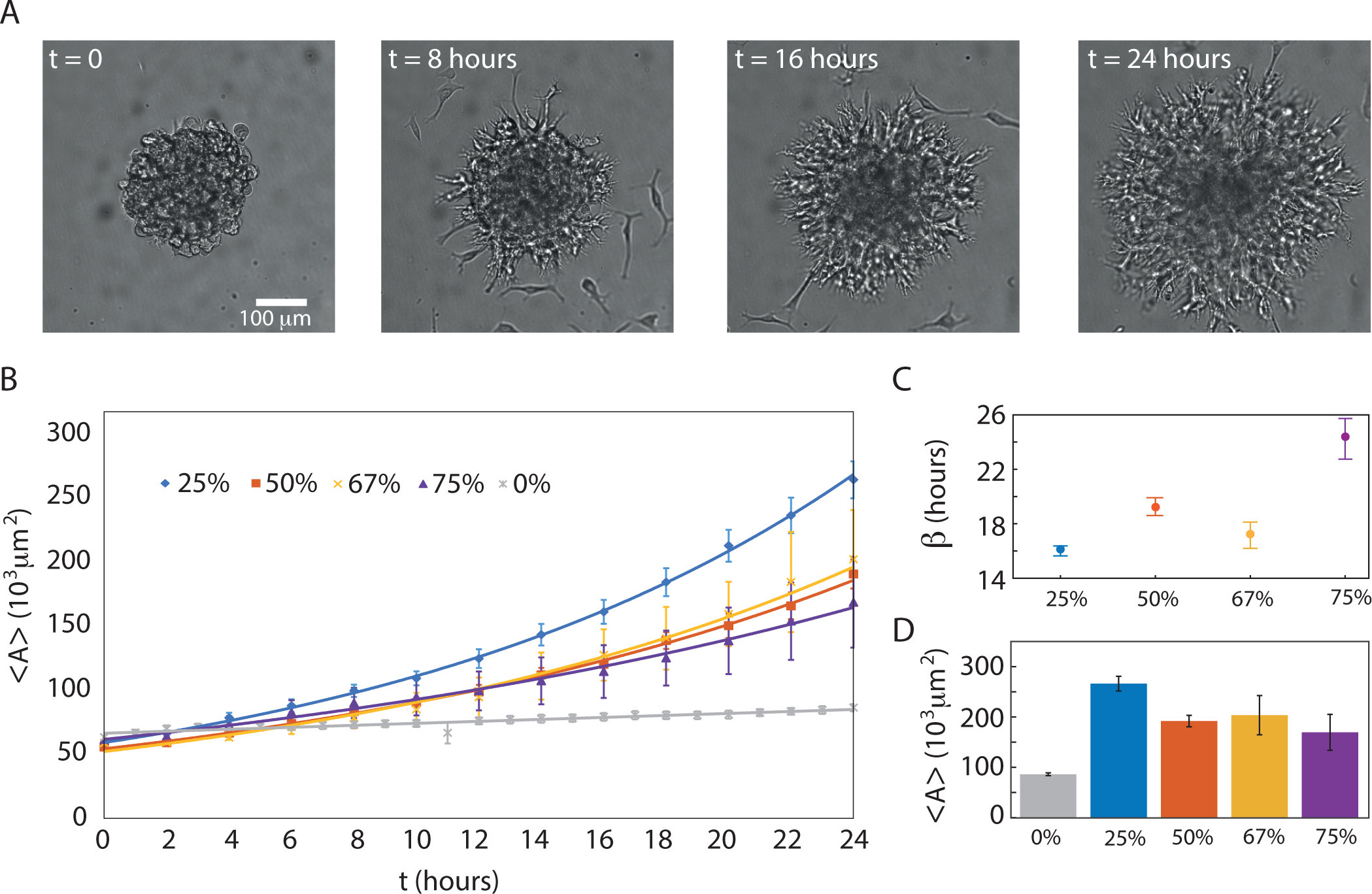
Invasion of brain cancer cells in ECM. **A)** Examples of migration/invasion of cells right after the transfer to ECM (50%) at time *t* = 0 and then after 8 hours, 16 hours and 24 hours, respectively. The star-shaped invasion pattern as well as individual migrating cells are clearly visible when *t* ≠ 0. **B-C)** Spheroid averaged invaded areas, ⟨*A* ⟩, vs. *t* for 25% (blue diamonds, *N* = 9), 50% (red squares, *N* = 8), 67% (yellow crosses, *N* = 4), 75% (purple triangles, *N* = 4), and 0% (grey asterisks, *N* = 9). Error bars are SEM and the full lines are exponential fit to the data with the growth exponents, *β*, as in equation 1 and given in (C) together with 95% CI. **D)** ⟨*A*⟩ at *t* = 24 hours, which corresponds to the last data points in (B) and error bars are SEM.

### Extra-cellular matrix stiffening slows down invasion

We measured the time-dependent average invaded cross sectional area, ⟨*A* (*t*) ⟩, as given by equation 3. Figure 1B shows the development for various ECM concentrations ranging from 0% to 75% ECM concentration. Here an ECM concentration of 0% corresponds to growth in medium only (grey asterisks) with an average growth rate of 38 [33-44] µm/day (*N* = 9). Hence, we confirm the linear growth of tumor spheroids in fresh culture medium Blumlein et al. 2017, Thorsen et al. 1997, Amaral et al. 2017. In contrast, for the invading spheroids, we found an exponential dependence between cross-sectional area and *t* with the characteristic growth exponent, *β*,

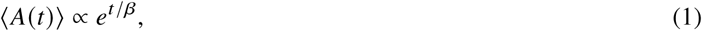

shown in figure 1C. To the best of our knowledge, an exponential growth of the spheroids’ cross-sectional area in ECM has not been reported before. It should be noted, however, that our findings of such fast expansion and invasion of the brain cancer spheroids for the first 24 hours after ECM addition are in agreement with previous reports of various cell lines invading type I collagen Kuo et al. 2017, Oraiopoulou et al. 2018. Disregarding spheroids in 67% ECM, *β* is responding linearly to the ECM concentration and, thus, the higher the ECM concentration, the slower the spheroids invade. However, for 67% ECM, we detect an abrupt decline of *β* compared to the slightly lower and higher ECM concentrations of 50% and 75%.

However, for all cases of non-zero ECM concentration, the spheroids’ final average areas, ⟨*A* (*t*) ⟩, are significantly larger than the 8.7 · 10^4^ µm^2^ spheroids in 0% ECM (grey bar) in figure 1D. This signifies that the resulting volume of an invading spheroid is larger than that of a non-invading - yet still proliferating - spheroid. The most pronounced difference is found for spheroids in 0% vs. 25% ECM (*p <* 0.001) while further increases in ECM concentration slows down invasion. For instance, for spheroids in 75% compared to 25% ECM, the invaded ⟨*A* (*t*) ⟩ drops by 1/3. Nevertheless, stiffening of ECM does not hinder invasion. While at slower pace, spheroids still invade and degrade the surrounding matrix. This is in accordance with findings for breast cancer cells in type I collagen Kuo et al. 2017, Gkretsi et al. 2017.

### Tumor cells migrate individually

After analyzing the collective invasive motility of the investigated brain cancer tumor spheroids, we tracked the cells that succeeded in detaching from the spheroid or invading cell collections. We followed migration through the ECM for several hours over hundreds of microns and with an algorithm that tracked the center-of-mass of the cells, we distinguished translocation from membrane dynamics. With this large number of data points, even low-probability events are captured.

In figure 2A, still images of spheroids are overlaid with examples of cell trajectories (colored lines) for the various ECM concentrations. Cell trajectories are highly diffusive, but in contrast to the invading cell collections, the migrating cells seem indifferent to ECM density at a first glance. This indifference is confirmed by the distributions of displacements, *d*, between two consecutive frames (5 minutes) as given by equation 5 and shown in figure 2B. Because of the heavy tails, these bar plots cannot be described by a Gaussian distribution as we would expect for random walk models Dieterich et al. 2008.

**Figure 2:**
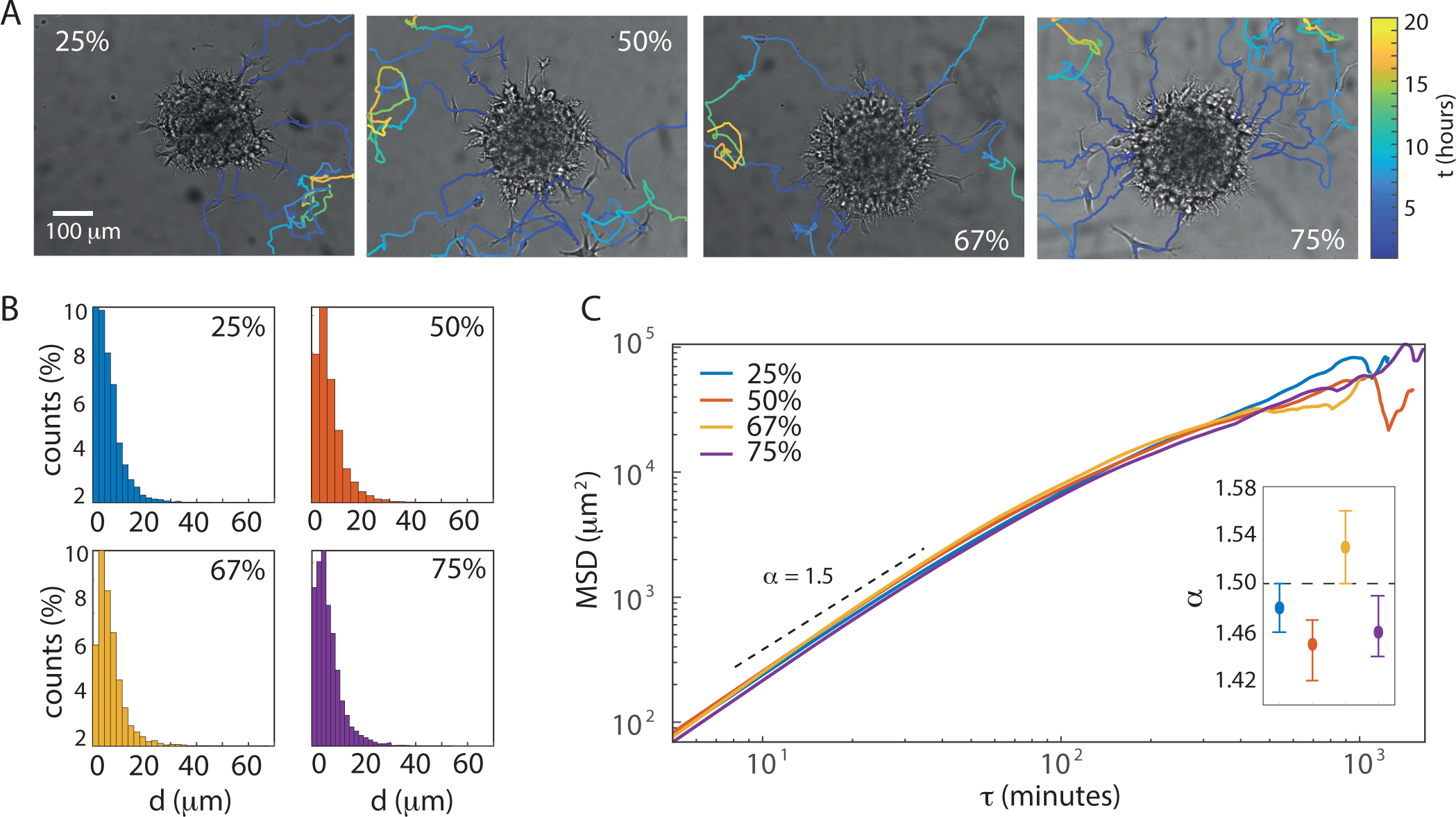
Cancer cell migration in various ECM concentrations. **A)** Examples of migrating cells’ trajectories from spheroids in various ECM concentrations. The color bar converts color to time, *t*. **B)** The distributions of displacements or steps, *d*, as given in equation 5. **C)** MSDs vs. the time delay, *τ*, for ECM concentrations of 25% (*N* = 223), 50% (*N* = 252), 67% (*N* = 123), and 75% (*N* = 108). The first 25% of each MSDs were fitted to equation 2 to obtain the anomalous exponents, *α*, which are given in the inset where error bars corresponds to 95% confidence intervals. As a guide to the eye, a dashed line with *α* = 1.5 has been inserted.

#### Super-diffusive migration

To further rectify this, we found the ensemble-averaged mean-squared displacement (MSD) as calculated in equation 6 and shown for the cell ensembles in 25% (blue), 50% (red), 67% (yellow), and 75% (purple) ECM concentrations in figure 2C. For completeness, mean-squared displacements of the individual cell trajectories for all ECM concentrations can be found in Supplementary figure S1.

The MSDs show anomalous diffusion of the migrating cells characterized by a power law dependence of the time delay, *τ*:

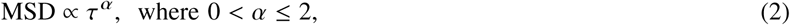

where *α* is the corresponding anomalous exponent. For normal diffusive motion *α* = 1. In contrast, anomalous diffusion is either sub-diffusive (*α <* 1), super-diffusive (1 *< α <* 2) or ballistic (*α* = 2). As 1 *< α <* 2 for all concentrations (inset of figure 2C), the migrating cells are super-diffusing which is a sign of active motility and in agreement with prior findings e.g. Takagi et al. 2008, Huda et al. 2018. Thus, the investigated cells appear to overcome the steric hindrances, i.e. the added strength of the physical barrier from increased protein density and decreased pore sizes, imposed by the increasing concentration of ECM. This is very different from what has been found for lung and bladder cells in collagen Laforgue et al. 2015, Plou et al. 2018, where concentration increases in collagen caused the cell motility to drop from super to sub-diffusive motion.

Moreover, the MSDs are almost indistinguishable up until *τ* ∼ 200 minutes. For *α* of 25%, 50%, and 75%, the 95% confidence intervals overlap, meaning that the anomalous coefficients are indistinguishable (inset of figure 2C). In contrast, the 67% *α* is significantly larger than for the other ECM concentrations, hence, cells migrate even more super-diffusively with this viscosity. Therefore, *α* evolves non-linearly with ECM concentration. For even longer lag times, *τ >* 1000 min, the MSDs exhibit erratic and erroneous behaviour due to sparse data.

#### Migration speeds are independent of extra-cellular matrix stiffness

We then investigated how modulating matrix stiffness affects the instantaneous speeds, *v*, of the individual steps along the migration trajectories; as given in equation 8. Box plots of the distributions of *v* for the various ECM concentrations are given in figure 3A. From these results we concluded that the increased ECM stiffness has no considerable effect on *v* for the migrating brain cancer cells. As *v* is linearly proportional to the displacements, *d*, we find the same heavy tailed distributions as in figure 2B, which signifies that cells most often migrate in small steps but once in a while in large leaps. Similar *bursts* of motion have been observed before Lee et al. 2012, Parker et al. 2018 and, hence, both the frequency and the amplitude contribute to the overall migration speed of the cells.

**Figure 3:**
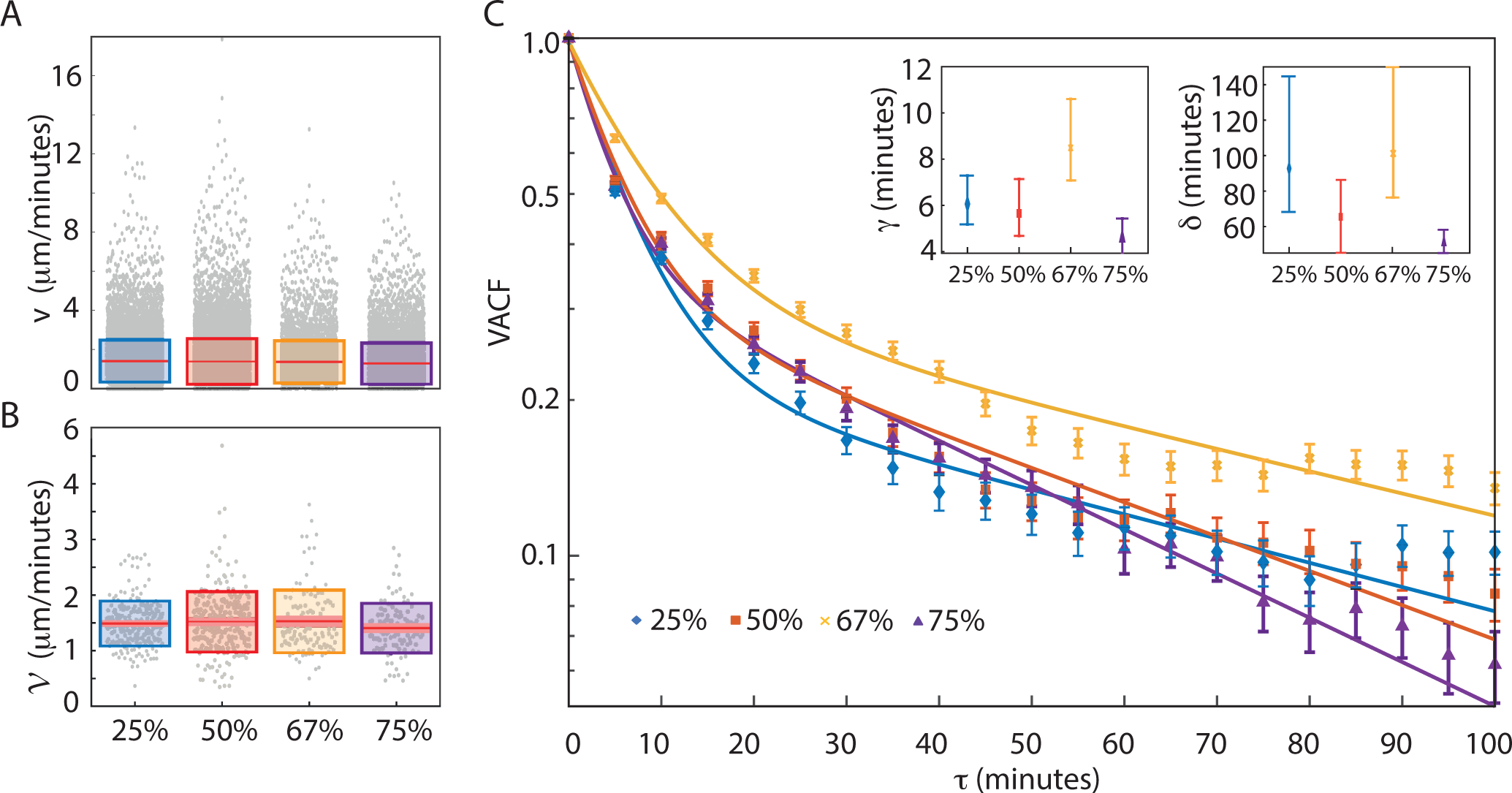
Effect of matrix stiffness on instantaneous speeds. **A)** Boxplot of instantaneous velocities, *v*, as defined in equation 8 for 25% (*N* = 15, 236), 50% (*N* = 17, 155), 67% (*N* = 6, 736), and 75% (*N* = 9, 452) ECM, where the red line signifies the median and the edges of the boxes from bottom to top are first (25%) and the third quartile (75%), respectively. **B)** Boxplot of the average migration speed, 𝒱, as defined in equation 9 for 25% (*N* = 223), 50% (*N* = 252), 67% (*N* = 123), and 75% (*N* = 108) ECM. Again, the distributions are indistinguishable despite the increasing stiffness of the matrix. **C)** Velocity auto-correlation functions, VACF, as defined in equation 10 for all ECM concentrations on semi-log scales. Points represent the data and error bars signifies one SEM. Full lines are the fits to a double exponential decay with characteristic decay times *γ* and *δ* for fast and slow decay, respectively. Insets show fitting parameters, *γ* and *δ*, for all ECM concentrations, where error bars represent 95% CI.

We also looked into the distributions of the migration speeds, *𝒱*, i.e., the average speed over the contour length of the trajectory, as defined in equation 9 and box plots of the distributions are shown in figure 3B. In contrast to the instantaneous speeds, the migration speeds are normally distributed with the following mean values: *𝒱* = (1.48 ± 0.40) µm/minutes (mean ± SD, *N* = 223), *𝒱* = (1.52 ± 0.54) µm/minutes (*N* = 252), *𝒱* = (1.53 ± 0.57) µm/minutes (*N* = 123), and *𝒱* = (1.41 ± 0.45) µm/minutes (*N* = 108) in the ECM concentration range from 25% to 75%, respectively. Thus, as verified by a Student’s *t*-test there are no significant differences between the speed distributions. Hence, the increased ECM rigidities cause no reductions of neither *v* nor *𝒱*, indicating a mechanism that allows for overcoming the increased tension of the surrounding matrix. As increasing ECM concentration does not only augment stiffness but also adhesion ligand density and decreases matrix porosity, the migration speed of cells in 3D ECMs is a result of the balance between stiffness, ligand density, proteolysis of the matrix and steric hindrances Zaman et al. 2006. For instance, in response to increasing ECM concentration pancreatic cancer has been shown to upregulate the activity of matrix metalloproteinases Haage and Schneider 2014, which are enzymes known to amplify cancer cell migrationValastyan and Weinberg 2011. Our results are in favor of a similar behavior, where migrating brain cancer cells boost their proteolytic activity to overcome the effects of increased matrix rigidity and adhesion as well as reduced pore sizes.

#### Velocity auto-correlation is a double exponential

To further investigate the nature of the motility, we found the velocity auto-correlation function (VACF) from equation 10 as presented in figure 3C. In general, the mean VACF is - strictly speaking - positive for all ECM concentrations, even at longer time scales. This behavior is consistent with a super-diffusive regime of motion where cells actively project forward in a direction dependent on the previous step. Ergo, the direction of motion is conserved in most cases. Although the correlation decays with time, the cells do not forget in which direction they were headed. Even for lag times up to *τ <* 100 minutes, velocities stay correlated. This memory effect is most pronounced for spheroids in 67% ECM and the least for those in 75% ECM which indicates that the increased matrix stiffness weakens the cells’ ability to keep a consistent direction of motion.

Mammalian cells’ motility has been studied for decades and most classic models are based on the Langevin equation and paths are described as Ornstein-Uhlenbeck processes of the persistent random walk (PRW). Here the MSD depends on diffusion, dimensionality, and the so-called persistence time, *τ*_*p*_, i.e. the time that the cell moves persistently in one direction Metzler and Klafter 2000, as given in equation 12. This model has been found to describe motions of e.g. fibroblasts Dunn and Brown 1987, Gail and Boone 1970, lung epithelial cells Wright et al. 2008, and neutrophils Parkhurst and Saltzman 1992. The Ornstein-Uhlenbeck process from equation 12 predicts a fast exponential relaxation Selmeczi et al. 2008. In contrast, we found a slower decrease of the VACF for long *τ*. The decays of the experimentally observed VACFs are, therefore, characterized by two characteristic relaxation times, *γ* and *δ*, shown in the insets of figure 3C. As expected from the fast decay of 75% ECM (purple line), the *δ* is in this case significantly smaller than for 25% and 67% ECM. Such density-dependent decay suggests that augmented ECM density dampens cell motility. Similar double-exponential decay has been found previously in e.g. human fibrosarcoma cell migration in 2D and 3D Wu et al. 2014, and in 2D migration of amoebas Takagi et al. 2008 as well as normal human dermal fibroblasts and human keratinocytes Selmeczi et al. 2005, while a gradual transition from exponential to power-law decay was found in mammalian kidney cells Dieterich et al. 2008.

#### Cells migrate both with directionality and persistence

To further investigate the directionality revealed from the VACFs, we studied the angular change between successive steps, *θ*, as defined in equation 11. For a randomly walking cell, we would expect a flat distribution of *θ* values, as the cell orients randomly in all directions. However, figure 4A shows that the *θ* distributions are skewed towards small angles. Therefore, the migrating brain cancer cells exhibit high directional persistence of displacements in all ECM concentrations. Figure 4B shows the distribution of *θ*s at increasing time lag, *τ*, in the range from 25 to 100 minutes as opposed to *τ* = 5 minutes between successive steps. The high occurrence of small angles does not flatten with time, even at *τ* = 100 minutes. Instead, the motion of the cells shows persistence, which goes directly against the predictions of the PRW model. Furthermore, the probability of observing large angles between two steps increases with increasing time lag. This is indicative of the tendency of the migrating cells to do 180° turns and backtrack; a behavior that is more pronounced in the higher ECM concentrations (67% and 75%), which is in accordance with other observations Wu et al. 2014, Geiger et al. 2019, Wu et al. 2015. A possible explanation for this is that cells - due to high adhesion and small ECM pore sizes - turn around and follow the beaten micro-tracks in the ECM back towards the spheroid. This asymmetric motility is described by the anisotropic persistence random walk (APRW) model, which encompasses both the high degree of heterogeneity between cells and the anisotropic movements Wu et al. 2015.

**Figure 4:**
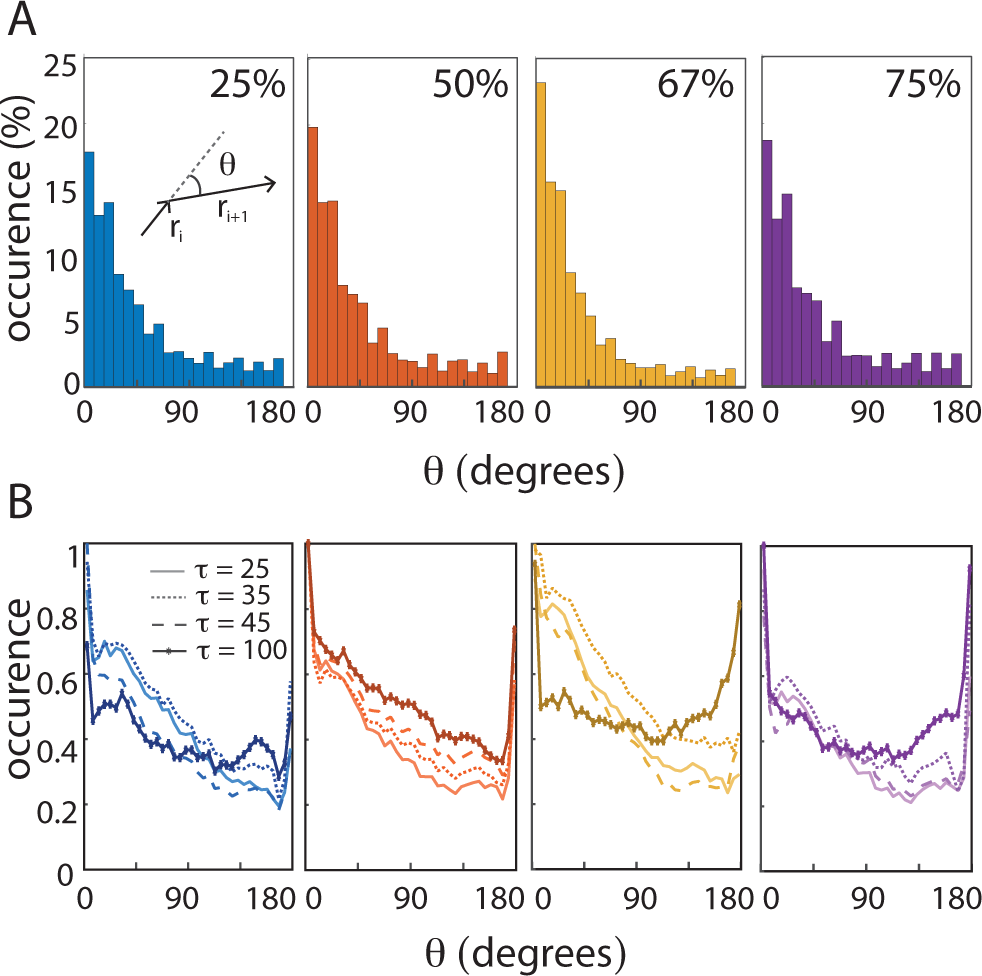
Directionality in increased ECM stiffness. **A)** Distributions of angles, *θ*, between two consecutive (*τ* = 5 minutes) steps, **r**_*i*_ and **r** _*i+*1_, as illustrated in the inset and defined by equation 11. The distributions for 25% (*N* = 14, 630), 50% (*N* = 15, 739), 67% (*N* = 6, 363), and 75% (*N* = 8, 719) ECM all show that the majority of *θ*s fall within |*θ*| *<* 90°. **B)** Angle distributions, *θ*, for different time delays, *τ*, from 25 to 100 minutes, see legend. As the counts vary for different *τ*’s, occurrences were normalized with respect to the counts for *θ* = 0.

#### Identifying super-spreaders

Based on these indications, we followed the train of thoughts of Wu et al. 2015 to fit the MSDs of individual migrating cells along their trajectory eigenvectors, primary (**p**) and non-primary (**np**), with the APRW model from equation 14. For each of the migrating cells’ trajectories, this model estimates the persistence time, *τ*_*p*_, along the primary axis of migration, **p**. The distributions of *τ*_*p*_ shown in figure 5A signify stochastic spreading dynamics with long ranged displacements Hallatschek and Fisher 2014. It has previously been reported that metastatic cancer cells do not only move in a super-diffusive fashion, but also display movement patterns consistent with Lévy walks Codling et al. 2008, Huda et al. 2018. We, thus, considered both power law (Lévy), exponential (Brownian) as well as log-normal models, which have been found to fit the motion of T-cells within lymph nodes Fricke et al. 2016. We found that *τ*_*p*_ followed a log-normal distribution for all investigated ECM concentrations. Even more surprising is the revelation of two distinct populations. Using *k*-means clustering we identified populations that are log-normal distributed around a lower *τ*_*p*_ (lighter color) and another with higher *τ*_*p*_ (full color). These two distinct log-normal distributions are characterized by their mean values and the SD of log(*τ*_*p*_). Hence, we find the mean *τ*_*p*_ to be the exponential of the log-normal mean and the so-called multiplicative-SD relates to the SD as *e*^SD^. Based on this, we find for the low *τ*_*p*_ populations (full color) average values of (12×/3.4) minutes (means ×/*e*^SD^, *N* = 171), (18.0×/3.8) minutes (*N* = 187), (14.9×/3.3) minutes (*N* = 87), (15.9×/4.2) minutes (*N* = 84) for ECM concentrations of 25%, 50%, 67% and 75%, respectively. All means of *τ*_*p*_ for these populations are around a quarter of an hour. In contrast, for the populations with higher *τ*_*p*_ (transparent color), we find averages of (3,442 2.4) minutes (*N* = 52), (3,393 2.6) minutes (*N* = 61), (3,302 2.4) minutes (*N* = 34), and (4,954×/2.0) minutes (*N* = 23) for ECM concentrations of 25%, 50%, 67% and 75%, respectively. Hence, these cells spread faster with long *τ*_*p*_; actually much longer than the total observation time (∼2,000 minutes). Therefore, we termed this population of highly persistent cells *super-spreaders*.

**Figure 5:**
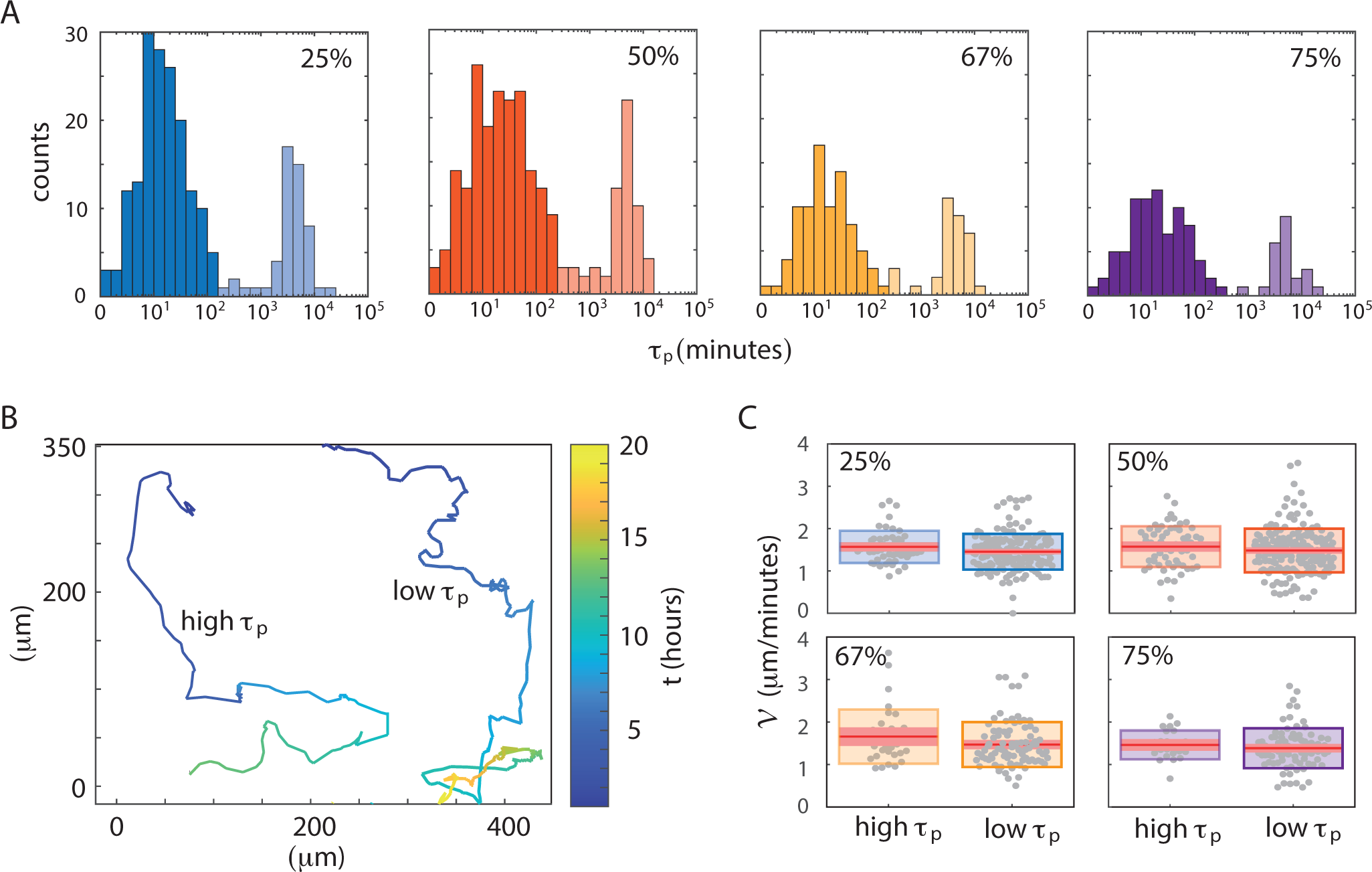
Identifying a distinct subpopulation of super-spreaders. **A)** Log-normal distributions of persistence times, *τ*_*p*_, for the various ECM concentrations. Using *k*-means clustering, we identified two distinct populations; one that is log-normal distributed around a lower *τ*_*p*_ (full color) and one around a higher *τ*_*p*_ (lighter color). **B)** Example trajectories from the majority population with low *τ*_*p*_ and the minority population with high *τ*_*p*_. Both examples are drawn from 75% ECM concentration and the color bar provides the conversion from color to time, *t*. **C)** Boxplot of the average migration speed, *𝒱*, from figure 3B redistributed with respect to high *τ*_*p*_ (lighter color) vs. low *τ*_*p*_ (full color) for all investigated ECM concentrations: 25% (*N* = 52, *N* = 171) for (high, low), 50% (*N* = 61, *N* = 187), 67% (*N* = 34, *N* = 87), and 75% (*N* = 23, *N* = 84).

An example trajectory drawn from the majority population of migrating cancer cells is given in figure 5B together with an example of a super-spreader cell’s trajectory. For completeness, we redistributed all tracks on the basis of high and low *τ*_*p*_ (Supplementary figure S2). It is obvious from these traces that the super-spreading cells very seldom stay within the field of view more than 10 hours after detachment from the spheroid body. As expected, this higher persistence of the super-spreaders is also reflected in the angle distributions. This is apparent from the subpopulation medians which become significantly different for all tested ECM concentrations between 25% to 67% when evaluated with a Wilcoxon rank-sum test (*p* ≪ 0.001). For 25% ECM the median *θ* is 28° and 34° for high *τ*_*p*_ and low *τ*_*p*_, respectively. Likewise for 50%, we found 26° vs. 31° and for 67% 21° vs. 27°. In contrast, for 75% ECM we found a less significant difference between the median *θ*s, 29° vs. 33° (*p* = 0.02), suggesting that the super-spreader motions are damped at very high ECM concentrations.

Following the surprising revelation of super-spreaders, we investigated whether this minority population was only characterized by more persistence i.e. less diffusive motion, or whether cells also moved faster along their path. Hence, we re-distributed the migration speeds, *𝒱*, as shown in figure 5C. However, the slight differences we found in average migrations speeds, *𝒱*, were non-significant (*p >* 0.05). So in conclusion, super-spreaders do not move faster - but more persistently.

## DISCUSSION

Spheroids are known to be very heterogeneous with an outer proliferating zone, an intermediate quiescent region with limited oxygen, nutrients, and metabolites Gole et al. 2009, Weiswald et al. 2010 as well as a necrotic core Vinci et al. 2012; exactly like solid *in vivo* tumors. Furthermore, when suspended in growth medium spheroids grow from the outer proliferating zone, but unlike many other species, the growth cannot be described by a logistic model, rather rates depend on population size Ben-Jacob et al. 2012, Korolev et al. 2014, Ron et al. 2003, Wallace and Guo 2013. When moved from serum-containing growth medium to ECM, our brain cancer spheroids invaded the surrounding matrix by long finger-like protrusions. This is a sign of collective migratory behavior where multicellular groups infiltrate the ECM through gained motility and retained cell-cell adhesion Jolly et al. 2015. These findings are paralleling earlier findings for similar spheroids (U87-MG) Vinci et al. 2012, Oraiopoulou et al. 2018 as well as for other glioblastoma spheroids (KNS42) Vinci et al. 2012, glioma spheroids (U-251-MG) Merz et al. 2015, and breast carcinoma spheroids Vinci et al. 2012, Daubriac et al. 2018. Therefore, this infiltrative nature makes it impossible to eliminate all cells in a host, thus leading to tumor recurrence.

From the acquired time-lapse sequences of invading spheroids, we found that the cross sectional area of the invading spheroid grows exponentially with time with no slowing of the growth rate within 24 hours. Nevertheless, a gradual stiffening of the ECM (Matrigel™) concentration slows down the invasion. Thus, the collective motility of cells in the finger-like protrusions responds to the polymer concentration in the matrix. An increased ECM concentration will necessarily also increase the concentration of individual ECM components such as growth factors (IGF, EGF, PDGF etc.) which activate and sustain cancer proliferation and growth Vukicevic et al. 1992, Kleinman and Martin 2005. Naively, one could therefore think that increased Matrigel™ concentration would augment the invasive capabilities of the spheroid. However, this augmentation is only observed when comparing ECM-embedded spheroids to spheroids in growth medium alone (0% ECM). In contrast, increasing the ECM concentration from 25% to 75% decreased spheroid invasion. We propose mechanosensing by the cancerous cells as a possible explanation for this. Several studies have shown that increased matrix stiffness triggers the mechanosensitive transcriptional activators YAP and TAZ which in turn activate e.g. proliferation, differentiation and metastasis in the tumor Jaalouk and Lammerding 2009, Dupont et al. 2011, Park et al. 2019, Rice et al. 2017. However, increased matrix stiffness is accompanied by a decrease in matrix porosity which limits the ability of cell collections to move through the matrix Anguiano et al. 2017. Like so many other functions in the tumor Zanconato et al. 2019, these mechanical cues might also be conveyed through YAP/TAZ and favor the full transformation into individual migrating cells rather than invasive cell collections. Mechanical feedback guides collective force generation Consequently, the mechanical stiffness of the tumor microenvironment not only triggers spheroid invasion, but also regulates it effectively through mechanical feedback Mark et al. 2020.

We further detected how individual cells detached from the spheroid - or the neighboring cells in the protrusion - to start individual migration through the ECM. This is indicative of a complete epithelial to mesenchymal transition, EMT, of the cells associated with weakening of the intercellular adhesion, loss of polarity, and increased motility Hu et al. 2016, Iwadate 2016. The cells were observed to first extend their pseudopodia, then displace the cell body followed by parallel retraction of pseudopodia. This behavior has been observed before in migrating glioblastoma cells Parker et al. 2018 and in the highly invasive human breast carcinoma cells (MDA-MB-231) Lee et al. 2012. Particularly, bursts of speed were pronounced when cells escaped from areas crowded by non-transformed breast epithelium cells (MCF10A). Moreover it was suggested that such fast movements contribute to the overall migration speed of each cell and they are the reason for their effective migration Parker et al. 2018.

Surprisingly, we found that glioblastoma cells exhibit similar migratory behaviors in all tested ECM concentrations. In contrast, if mechanosensing causes a shift from invasion towards migration as hypothesized above, we would expect an increase in the overall migration speeds or diffusion constants with ECM concentration. The notion that increased stiffness increases cancer cell migration due to YAP/TAZ activation supports this argument. However, Matrigel™ has been found to play a dual role: While the presence of Matrigel™ in collagen gels promoted migration of lung cancer cells (H1299) at low concentration, it hindered migration at higher concentrations, probably due to increased adhesion-ligand concentration Anguiano et al. 2017. Thus, our hypothesis - that increased ECM concentration favors increased single cell migration - might be counteracted by increased adhesion-ligand concentration and result in no apparent ECM concentration dependence Geiger et al. 2019. Moreover, earlier studies suggest that proteolysis serves as an important additional mechanism for migration of cancer cells, allowing them to move through the matrix - no matter how dense it is Zaman et al. 2006, Valastyan and Weinberg 2011, Koh et al. 2018. Further, it has been established that during collective cell invasion, the leading edge cells carve paths for the remaining cells by proteolytic remodeling of the ECM Khalil and Friedl 2010, Trepat et al. 2009. For example for fibrosarcoma cells (HT-1080) invading 3D collagen, leader cells utilize proteolysis by a membrane-bound collagenase (MT1-MMP) to generate tracks Wolf et al. 2007. Although we found the average anomalous exponent to be *α* ∼ 1.5, we observed both Brownian-like single cell trajectories with *α* ∼ 1, as well as cells with *α >* 1.5 at all ECM concentrations. This is indicative of a similar leader/follower relationship for migrating cells: leader cells with Brownian-like diffusion proteolytically generate paths for subsequent cells to follow. Because paths have been generated for them in advance, follower cells exhibit super-diffusive motion with high directionality and persistence. Further, it has been shown that collagenase activity in pancreatic cancer cells (Panc-1) increases with matrix stiffness Haage and Schneider 2014. This indicates that leader cells have mechanosensing abilities that cause them to increase proteolytic activity in order to overcome increased matrix stiffness, thus apparently rendering migration of U87-MG glioblastoma cells ECM-concentration independent. Nevertheless, we did observe slight concentration dependencies, such as decreased VACF decay times and increased probability of backtracking at long time scales in high ECM concentrations.

Any Brownian random walk model of cell migration would predict a uniform distribution of turning angles between successive steps, while the PRW model predicts a peaked distribution that flattens at time scales longer than the persistence time. In contrast to both these random walk models, we found that the probability of complete 180° turnarounds in the 3D matrix increased with time. This result indicates that the probability of observing cells backtracking through the tunnels - formed by the cells during their initial exploration - increases at longer time scales. Given such anisotropic persistence, we fitted our results to the APRW model and extracted the according persistence time distributions for the primary directions of migration. We found long-range displacement dynamics interspersed between long periods of short-range steps, which is characteristic of Lévy walks. Furthermore, a recent study proved that the motion of metastatic cancer cell lines performed Lévy walks, while their non-metastatic counterparts did not Huda et al. 2018. As Lévy walks also have uniformly distributed turning angles, the arguments for describing the motion of migrating glioblastoma cells as a Lévy walk were, hence, very compelling. However, our persistence time distributions did not follow a power-law. Instead we - unexpectedly - found the persistence time distributions to be log-normal and bimodal. This is suggestive of the existence of two subpopulations of which one is a population of *super-spreaders* with boosted persistence. Such contradictory observations have been done previously, e.g., for immune T-cell motion, which has been identified as either log-normal (Fricke et al. 2016) or a Lévy walk Harris et al. 2012 depending on the tissue they migrated in. Moreover, evidence that individual persistent random walkers exhibit Lévy walk movement patterns at the ensemble level has been published in the case of single bacteria vs. swarms Ariel et al. 2015, Reynolds and Ouellette 2016. Evidently, the distinction between variants of PRWs and Lévy walks warrants further investigation Auger-Méthé et al. 2015, Svensson et al. 2018.

In summary, we analyzed cell migration from glioblastoma spheroids embedded in varying concentrations of ECM-like Matrigel™. While collective migration was partially inhibited by high-density gels, single cell migration was not. However, migration of single cells did become more anisotropic with increasing ECM density. Modeling the migration as an APRW, we identified a subpopulation of super-spreading cells with extraordinary directional persistence. Development of a model that captures all the behaviors we have observed - namely double-exponential decay of the VACF, highly directional and persistent migration, anisotropic behavior at high ECM density and the existence of super-spreading cells - would allow prediction of the motility mode of the migrating cells further from the spheroid. This would be an important step en route to a unifying model of cancer metastasis. Moreover, although the concept of intra-tumoral heterogeneity has been established years ago, it is still an area of research the receives much attention. Our work presents yet another aspect of between-cell tumor heterogeneity with high clinical relevance since targeting the population of super-spreading cells would be the most efficient methodology for prevention of metastasis and relapse after surgery. Future studies will have to uncover whether our observations represent actual differences between cells within the tumor spheroids or whether they arise in response to different local environments.

## MATERIALS AND METHODS

### Cell culture

The glioblastoma multiforme cell line U87-MG was cultured under standard conditions (37°C, 5% CO_2_, 95% humidity) in Dulbecco’s Modified Eagle Medium (DMEM) supplemented with 10% Fetal Bovine Serum (FBS) and 1% penicillin-streptomycin; all provided from Gibco (Gibco, Life Technologies Ltd. Paysley, UK).

#### Gravitation-assisted tumor spheroid formation

For the formation of spheroids, 650 cells (3250 cells/mL) were seeded in ultra-low attachment round-bottomed 96-well plates (Corning B.V. Life Sciences, Amsterdam, The Netherlands) with 200 µL DMEM and incubated for 4 days Niora et al. 2020.

#### Spheroid invasion/migration

Our invasion assay was based on classic protocols described by Vinci et al. 2012 and Richards et al. 2018. Matrigel™ (Corning B.V. Life Sciences, Amsterdam, Netherlands) was thawed on ice overnight at 4°C. On the day of experimentation, the round-bottomed well plates with fully-formed spheroids were placed on ice to avoid untimely Matrigel™ solidification. Then, medium was removed from the wells and replaced with ice-cold Matrigel™ (using pipette tips kept at −20°C) to the desired final concentration (v/v) of either 25%, 50%, 67%, or 75%. When interchanging medium with Matrigel™ we took care to keep the spheroids centered in the bottom of the well and to remove all bubbles. Afterwards, plates were incubated for 1 hour for solidification of the Matrigel™ and lastly 100 µL DMEM was added on top of each well to a final volume of 300 µL.

### Time-lapse imaging

96-well plates were mounted on an inverted Juli Stage Real-Time cell history recorder (NanoEntek, Guro-gu, Seoul, Korea) with bright-field (10× objective) placed inside a cell incubator (37°C, 5% CO_2_, 95% humidity). The focus of the microscope was set manually for each well and we used the built-in software to acquire time-lapses with a frame rate of 1 every 5 minutes, exposure time of 170 ms, and field of view of approximately (850×640) µm^2^ with a resolution of 440 nm/pixel. For 24 or 31 hours of imaging, 50 µL of DMEM was added to the wells during experimentation to prevent dehydration of wells. Please, note that our imaging modality results in a 2D approximation of the 3D migration as discussed in detail by Wu et al. 2014.

### Image analysis

#### Spheroid invasion

To quantify spheroid invasion we used the Fiji - ImageJ Schindelin et al. 2012 macro tool *Analyze Spheroid Cell Invasion In 3D Matrix* to obtain the projected area of the main spheroid body Baecker 2012. From the resulting cross sectional area, *A*, we found the average area of all spheroids to be:

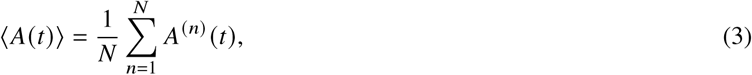

where *A*^(*n*)^ (*t*) is the area of the *n*-th tumor spheroid at time *t* and *N* is the total number of spheroids.

#### Image pre-processing

We used the open-source software ilastik, which enables automated - albeit supervised by the user - segmentation of spheroids and individual cells Sommer et al. 2011. For training of the algorithm, we used a representative stack of 50 consecutive images from 0 to 250 minutes to minimize computation time. Please note that some manual track-editing was needed for our data because of over-segmentation, i.e., one cell is detected as two, and under-segmentation (vice versa). We also had to remove data points from objects (of size similar to cells) that mistakenly had been detected as cells. All these errors were filtered manually to ensure meaningful results.

#### Single cell migration

For automated single particle tracking we used the *TrackMate* plugin for Fiji - ImageJ, which consists of a *spot detection* followed by a *particle tracking* procedure. Since our cells expand over a rather large area in the segmented image (⩾ 20 pixels ∼ 9 µm), we used the *Downsample LoG* detector Tinevez et al. 2017 with the down sample factor set to 4, the estimated blob diameter to 50 µm, and a cell/background threshold value of 1 appropriate for our binary images. With these settings, the spot detection algorithm detected all centers of mass of migrating cells as well as cells on the spheroid surface. Once spot detection is completed, cells were tracked in every frame of the image stack using the *The Simple Linear Assignment Problem* algorithm since - upon visual inspection of the images - mitosis events were rare within our time frame. For post-processing, we used the *TrackScheme* module to edit the cell trajectories and we ended a track in the following cases: i) if a cell moved out of the field of view (even if it returned later), ii) if cells never really detached from the spheroid before returning, and iii) after cell division if it occurred Mitchell et al. 2018.

### Data analysis

#### Displacement analysis

An ensemble of particles undergoing Brownian motion is distributed as Bian et al. 2016:

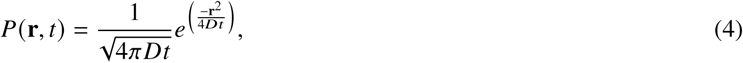

where *D* is the diffusion coefficient and *t* is the time step. In the *n*-th cell trajectory we define a position vector in the projected plane as: **r**^(*n*)^ (*t*). The length of the vector is the displacement, *d*:

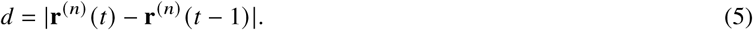

#### Mean squared displacement analysis

The mean squared displacement (MSD) describes the displacement of cells by determining the ensemble-averaged distance that the ensemble of cells move in a given time interval, *τ*, which is designated as the lag or delay time Tarantino et al. 2014:

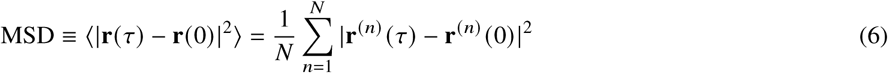

where *N* is the total number of trajectories for all measured spheroids, vector **r**^(*n*)^ (0) is the starting position of the *n*-th cell trajectory, and vector **r**^(*n*)^ (*τ*) is the position of the *n*-th cell after time lag *τ*.

The MSD of a random walk, i.e., normal diffusion, of cells in 3D scales linearly with time:

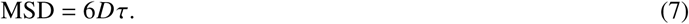

For finite trajectories, the small *τ*’s are weighted more in the average MSD than longer delays. For example, if a trajectory has *N* points, the delay, *τ*, that is equal to 1 time unit will have *N* − 1 points in the average, whereas the delay that corresponds to *N* time units will have only 1 point in the average Miura and Sladoje 2019. Therefore, the accuracy of the MSD becomes increasingly worse with *τ* as the variance depends on the number of trajectories available. Moreover, the curvature of the MSD is important when investigating whether there is active transport in the system under study. In the ensemble-averaged MSD, there are contributions from different kinds of motion which makes the curvature change over time thus increasing statistical uncertainty.

Therefore, it has been suggested that to limit the relative errors of the MSD, the plots should not be analyzed and fitted to any model further than 25% of the longest time lag. Furthermore, when fitting an MSD to a model, one should use proper weights, such as the inverse of the variance or standard deviation for each time lag, in order to give more weight to the parts of the MSD with the highest certainty Saxton 2007, Diaspro 2010. Both suggestions were adopted in this study.

#### Instantaneous speed distributions and migration speed

We defined the instantaneous speed, *v*, at the *i*-th step of a cell as the displacement in the projected *x,y*-plane per time step:

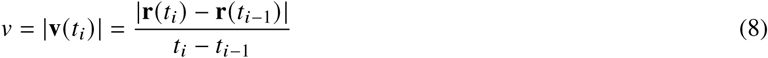

and the migration speed of a cell, *𝒱*, as the mean over the entire contour length of the trajectory from *i* = 1 to the total number of steps, *i* = *L*:

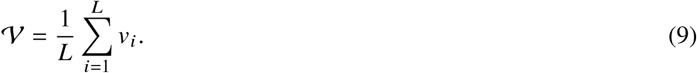

#### Motion persistence and directionality

We also investigated the velocity auto-correlation function (VACF) of the ensemble average of cell trajectories which is:

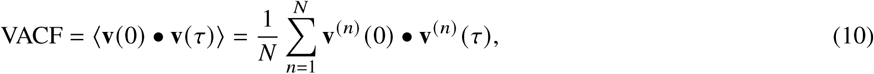

where the velocity at time *τ* is **v**^(*n*)^ (*τ*) of the *n*-th cell trajectory and the initial velocity is **v**^(*n*)^ (0). In general, the VACF diminishes with increased *τ*, as the cells’ velocities become uncorrelated due to interactions with the surrounding environment. Furthermore, the angle between the position vectors between two successive steps **r**(*t*_*i*_) and **r**(*t*_*i*−1_) fulfill the relation:

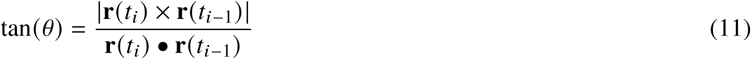

#### The Ornstein-Uhlenbeck process

The Ornstein-Uhlenbeck process describes a random walk model but with an added persistence (PRW) Selmeczi et al. 2005. This model of cell motility is derived from a stochastic differential equation describing the motion of a self-propelled cell. By introducing persistence, the MSD in 3D as given in equation 7 for a simple random walk is described by Fürth’s formula Campos et al. 2010:

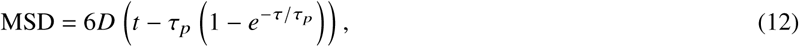

where *τ*_*p*_ is the persistence time of the motion, i.e., the time for which a certain velocity is retained by the system Selmeczi et al. 2008. Therefore, *τ*_*p*_ is the time before a cell changes direction. Equation 12 can be rewritten as:

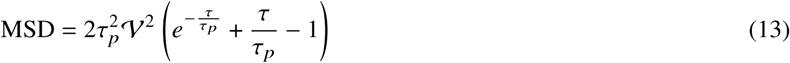

if we assume an exponential distribution of persistence times Potel and Mackay 1979. Furthermore, the Ornstein-Uhlenbeck velocity auto-correlation function, VACF, which can be derived by differentiating equation 12, is predicted to be a single-exponential decay.

#### Anisotropic persistent random walk model

We have adopted the anisotropic persistent random walk model (APRW) for the migration of cells in a 3D matrix Wu et al. 2014, 2015. In the APRW, the velocity of the migrating cells is assumed to be spatially anisotropic and display different persistence times on two orthogonal axes, namely the primary **p** migration axis and the non-primary **np** migration axis; these are found through singular vector decomposition of cell velocities.

The migration of the cells in the APRW model is described by the same parameters as in the PRW model, the persistent time *τ*_*p*_ and cell speed *𝒱*V, however they are different for each individual axis of motion. Therefore, for the primary axis they are denoted as (*𝒱*_*p*_, *τ*_*p*_) and for the non-primary they are (*𝒱*_*np*_, *τ*_*np*_). The MSDs in the primary direction, **p**, are given by:

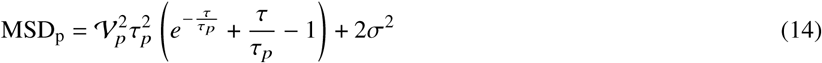

and in the non-primary orthogonal direction **np**:

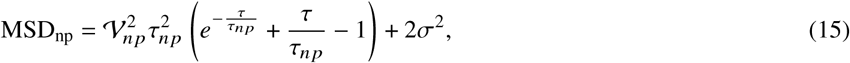

where 2*σ*^2^ is the error in the position of the cell and the total MSD for each cell is given by:

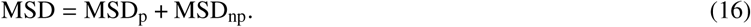

## AUTHOR CONTRIBUTIONS

AN did the spheroid imaging and single-particle tracking, MTS did supportive experiments, and LJ supervised the project. All authors discussed the results and contributed to the final manuscript and all authors read and approved the final manuscript.

## ACKNOWLEDGMENTS

LJ acknowledges funding from the Danish National Research Council (DNRF116) and AN thanks Kristoffer Laugesen, Younes Farhangibarooji, Guillermo S. Moreno Pescador, Mads Kasper von Borries and Ioannis Koutsoukidis for fruitful discussions.

## SUPPLEMENTARY MATERIAL

**Figure S1:**
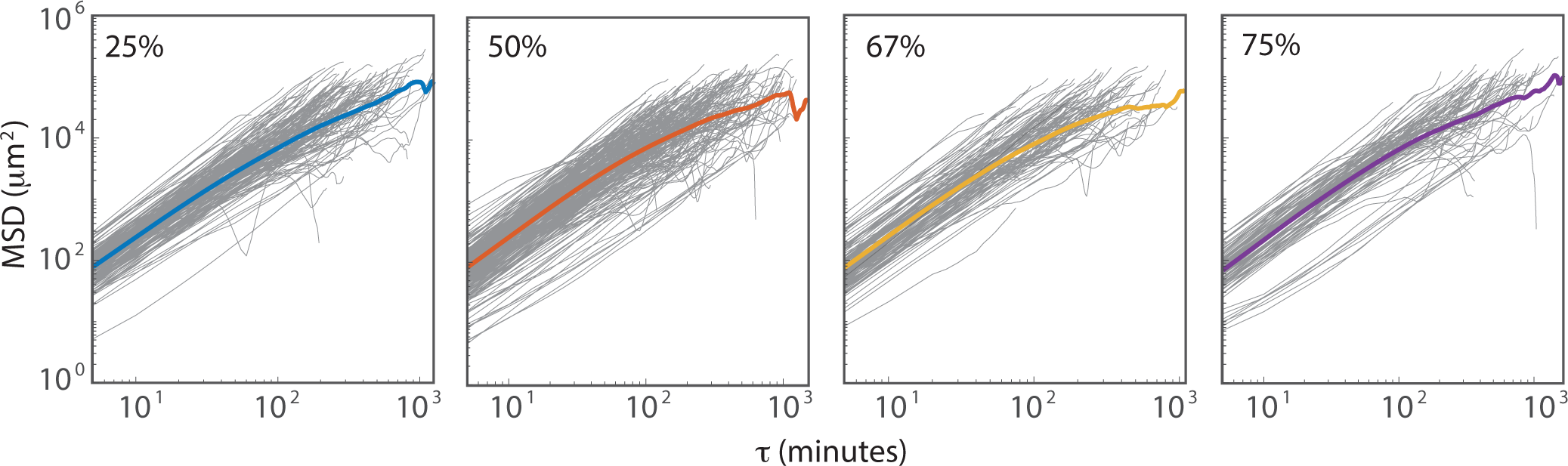
All mean-squared displacements of individual trajectories (light grey) and the mean MSD (thick colored line) at 25%, 50%, 67%, and 75% ECM concentration. The MSD is given in figure 2.

**Figure S2:**
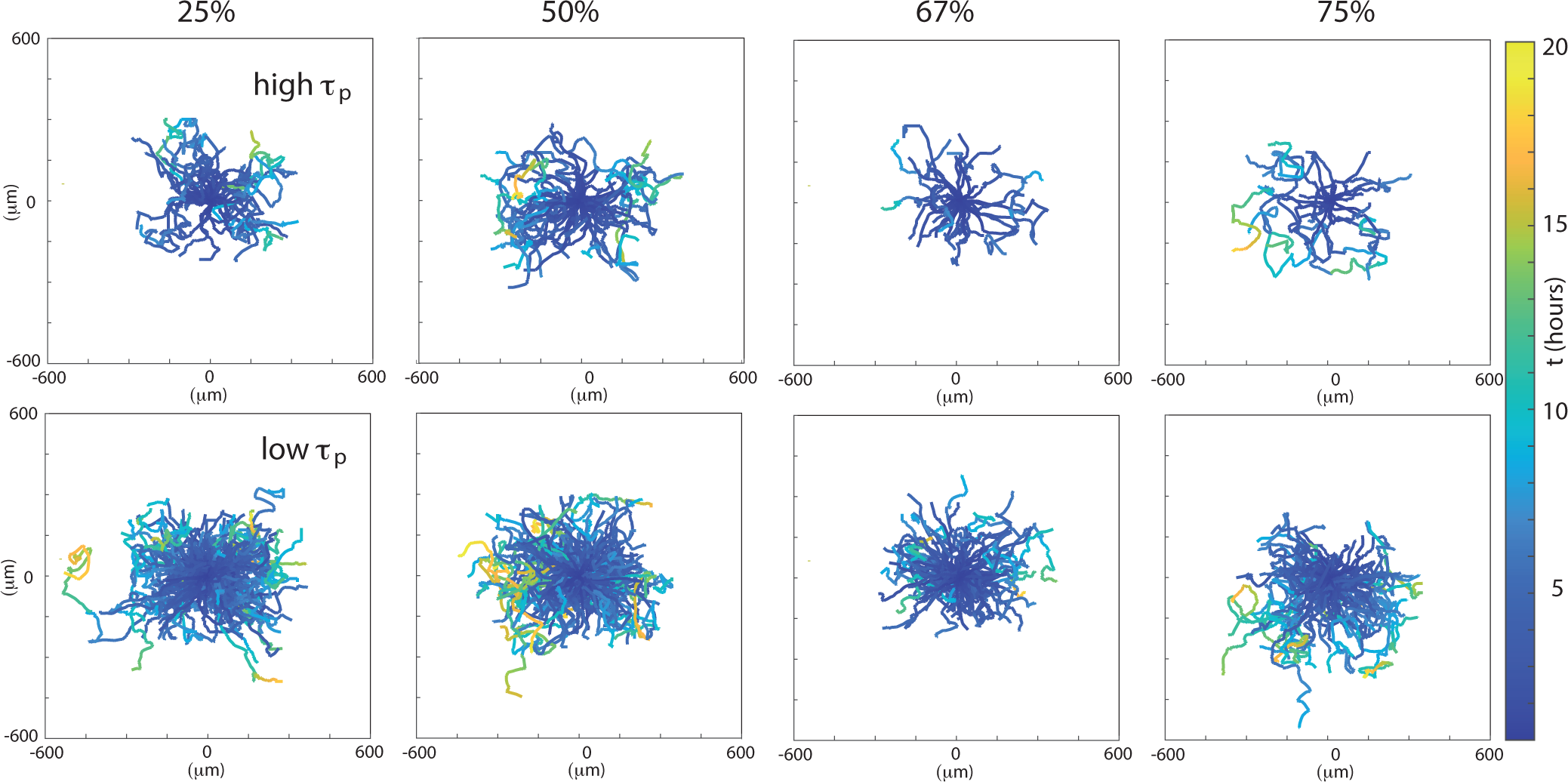
All trajectories for either high *τ*_*p*_ (upper panel) and low *τ*_*p*_ (lower panels) for 25%, 50%, 67%, and 75% ECM concentration, respectively. The redistribution of trajectories in respect to high/low *τ*_*p*_ is done on the basis of figure 5.

